# Event-Related Potentials of Single Sided Deaf Cochlear Implant Users – Using a Semantic Oddball Paradigm in Noise

**DOI:** 10.1101/2022.06.30.498355

**Authors:** Marcus Voola, Andre Wedekind, An T. Nguyen, Welber Marinovic, Gunesh Rajan, Dayse Tavora-Vieira

## Abstract

**Objective:** In individuals with single sided deafness (SSD), which is characterised by a profound hearing loss in one ear and normal hearing in the contralateral ear, binaural input is no longer present. A cochlear implant (CI) is the only way to restore functional hearing in the profoundly deaf ear, with previous literature demonstrating improvements in speech in noise intelligibility with the provision of a CI. However, we currently have a limited understanding of the neural processes involved (e.g., how the brain integrates the electrical signal produced by the CI with the acoustic signal produced by the normal hearing ear) and how the modulation of these processes with CI contributes to improved speech in noise intelligibility. Using a semantic oddball paradigm presented in the presence of background noise, this study aims to investigate how the provision of CI impacts speech in noise perception of SSD CI users.

**Method:** High density electroencephalography (EEG) from twelve SSD-CI participants was recorded whilst they completed a semantic acoustic oddball task. All participants completed the oddball task in three different free field conditions with the speech and noise coming from different speakers. The three tasks were 1) with the CI-On in background noise, 2) with the CI-Off in background noise and 3) with the CI-On without background noise (Control). We examined task-performance (RT, subjective listening effort, and accuracy) and measured N2N4 and P3b event-related brain potentials (ERPs) linked to the discrimination and evaluation of task relevant stimuli. Speech in noise and sound localisation abilities was also measured.

**Results:** Reaction time was significantly different between all tasks with CI-On (*M*(*SE*) = 809(39.9) ms) having faster RTs than CI-Off (*M*(*SE*) = 845(39.9) ms) and Control (*M*(*SE*) = 785(39.9) ms) being the fastest condition. The Control condition exhibited a significantly shorter N2N4 and P3b area latency when compared to the other two conditions. However, despite these differences noticed in RTs and area latency, we observed similar results between all three conditions for N2N4 and P3b difference area.

**Conclusion:** The inconsistency between the behavioural and neural results suggest that EEG may not be a reliable measure of cognitive effort. This rationale is further supported by the different explanations used in past studies to explain N2N4 and P3b effects. Future studies should look to alternative measures of auditory processing (e.g., pupillometry) to get a deeper understanding of the underlying auditory processes that facilitate speech in noise intelligibility.

## Introduction

Single sided deafness (SSD) is characterised by a profound hearing loss in one ear and normal hearing in the contralateral ear [Friedmann et al., 2016]. Unlike individuals with a bilateral hearing loss, SSD individuals can rely on the normal hearing ear (NHE) to understand speech in quiet, thereby reducing the impact that the hearing loss has on quality of life [Voola and Távora-Viera, 2021]. However, in noisy environments, speech intelligibility of SSD individuals decreases significantly when compared to normal hearing individuals [Van de Heyning et al., 2008;Williges et al., 2019;Körtje et al., 2022]. Speech intelligibility in noise is facilitated through the binaural squelch and binaural summation effects, both of which rely on similar inputs from both ears [Ma et al., 2016]. A cochlear implant (CI) is the only treatment option that has the potential to restore binaural hearing, thereby providing access to the advantages of bilateral hearing which in turn can improve speech understanding of SSD individuals in background noise.

A CI has the potential to restore hearing by directly stimulating the auditory nerve in the impaired ear via electrical signals [Drennan and Rubinstein, 2008]. Sound transmission through the CI is degraded and does not fully encapsulate all the spectral information that the NHE provides [Drennan and Rubinstein, 2008]. Despite this limitation, a CI for SSD individuals can improve speech intelligibility in noise and localisation ability [Távora-Vieira et al., 2015;2016;Dorbeau et al., 2018;Galvin et al., 2019;Williges et al., 2019;Wedekind et al., 2020]. These improvements highlight that the brain is capable of understanding both the acoustic signal from the NHE and the degraded electrical signal from the CI. However, it is not well understood how the underlying neural process operate to improve speech in noise intelligibility. One method to understand how the neural process are operating is by using EEG to examine event related potentials (ERP) evoked by the presentation of acoustic stimuli. Together, these measurements can provide an insight into the cortical processing of auditory stimuli.

Auditory ERPs have been used in the past to measure the neural processing of auditory information in CI users. ERPs are characterised by a series of deflections with a fixed time-course. Scalp-distribution and amplitude differences in these deflections provide an insight into the different stages of auditory processing [Light et al., 2010]. Auditory ERPs can be elicited by an oddball paradigm, this consists of a frequent (standard) and non-frequent (target) stimuli whereby participants are instructed to indicate when they hear the target stimuli. The target stimuli can differ from the standard stimuli in multiple ways such as differences in physical properties (e.g., frequency, intensity) or semantic qualities (e.g., living vs. non-living words) [Polich, 1985;Polich et al., 1990]. However, understanding how higher order processing facilitates speech in noise understanding in SSD CI users has yet to be investigated.

Higher order neural processing that involves discrimination and evaluation of stimuli is reflected through changes in a fronto-central negativity (N2) and parietal positivity (P3b). The N2 deflection occurs within a latency range of 200 – 350 ms after stimulus onset and is enhanced upon the presentation of the target stimuli thereby reflecting the process of discrimination [Lau et al., 2008]. As task difficulty increases in complexity, a delayed peak latency is observed which is thought to represent the difficulty in discriminating the stimuli from stored mental representation [Näätänen and Picton, 1986]. For more complex tasks, such as those involving the discrimination based on semantic meaning rather than pure tone differentiation, the N2 peak latency can be delayed to around 400 ms, resulting in the peak to be labelled as the N4. This delay in peak latency is attributed to the additional time needed for individuals to fully retrieve the words meaning from their stored mental lexicon. However, differentiating the N2 from the N4 is challenging with many studies reporting difficulties in distinguishing the two [Deacon et al., 1991;van den Brink et al., 2001;Finke et al., 2016]. As such, to avoid confusion and to follow in line with previous studies, the second negativity of the ERP waveform will be referred to as the N2N4.

The process of stimulus evaluation and categorising is represented via a parietally distributed positive deflection occurring at a latency of 300 – 600 ms, referred to as the P3b [Polich, 2007]. The P3b is thought to represent the process of decision making, whereby the presentation of a stimulus triggers the activation of stimulus-response links [Verleger et al., 2014]. Stimuli that are more demanding (i.e., differentiating stimuli based on meaning) have been identified to elicit a smaller P3b amplitude and delayed latency [Polich, 1985;1986;Johnson, 1988;Verleger, 1997;Comerchero and Polich, 1999]. Additionally, past studies have identified that more involved tasks result in larger reaction time (RT) which provides support for the decision-making hypothesis.

Literature focusing on the higher order processing of CI users in background noise is limited. Soshi et al (2014) investigated how the P3b is affected by noise in the CI population by presenting /ga/ and /ba/ syllables. It was identified that only good performing CI users (speech perception in noise score is greater than 66%) were able to elicit a P3b in noise, suggesting that the speech perception scores in CI users is positively correlated to their P3b amplitude [Soshi et al., 2014]. In 2016, Finke et al. (2016) built upon the work of Soshi et al by instructing subjects to differentiate words as either living or non-living entities in the presence of background noise when presented in free field. Using a more complex stimuli in the form of semantic differentiation, rather than just differentiating based on physical properties provides a firmer representation of how the higher order processes of CI users are working in everyday life. Compared to the normal hearing control, CI users exhibited delayed N2N4 and P3b latency, increased RT which may be attributed to the mismatch between the limited CI input and the stored mental representations [Finke et al., 2016].

Given the unique hearing loss of SSD CI recipients (normal hearing in one ear and a profound hearing loss in the contralateral ear) this provides a unique opportunity to isolate the impact of the CI by employing a within-subject designed experiment. Finke et al (2016) and Wedekind et al (2021) both identified that direct stimulation of the CI requires greater processing effort (as indicated by delayed RTs) when compared to stimulation of the NHE alone in SSD CI users [Finke et al., 2016;Wedekind et al., 2021]. Whilst these studies do provide a foundation for understanding CI processing, they do not address how the electrical signal from the CI and acoustic signal from the NHE is integrated at a cortical level to provide binaural benefit. As such, in a previous study conducted by our team, we presented semantic stimuli to SSD CI users with the aim to identify how the higher order neural processing differs with and without the CI in free field. We found clear evidence that in free field the brain is processing the input from both ears when the CI is on as indicated by a significantly enhanced P2 amplitude. However, the behavioural results indicated that the addition of the CI lead to greater uncertainty (larger RT variability) and delayed RT, which lead us to believe that the speech in quiet task was not well set up to assess binaural hearing. This rationale is further supported by the fact that the task used did not evaluate the binaural squelch effect (an advantage of binaural hearing), thereby the normal hearing ear was able to evaluate and discriminate the speech in quiet [Voola et al., 2022]. As such, this study was designed to build up on the findings of Voola et al 2022 by incorporating background noise into the semantic oddball task, thereby aiming to investigate how the CI impacts speech in noise perception of SSD CI users. We hypothesize that in the CI-Off condition SSD CI users will have poorer speech in noise discrimination, which will be reflected by delayed RT and smaller and delayed N2N4 and P3b effects. SSD CI users will perform better in the CI-On condition. Using more complex variations of the oddball paradigm (i.e., in noisy environments) may provide a more thorough understanding of the underlying neural processes that facilitate higher order processing in SSD CI user.

## Materials and Method

### Participants

Twelve SSD CI participants were recruited from the Fiona Stanley Hospital audiology department. Three participants were also part of previous study of ours Voola et al 2022. All adult participants (> 18 years) were required to have normal hearing in one ear, which was defined as having a four-frequency average (250 Hz, 1 kHz, 2 kHz, 4 kHz) hearing loss less than or equal to 20 dB HL. In the contralateral ear, all SSD CI participants have been using a MED-EL cochlear implant for at least one year. Participants gave written informed consent prior to participating in the experiment. Ethics approval was obtained from the South Metropolitan Health Ethics Committee (reference number: 335).

### Speech Perception in Noise

The Bamford-Kowal-Bench Adaptive Speech-In-Noise test was used to measure the speech in noise intelligibility of the SSD CI participants [Bench et al., 1979]. Each participant underwent the assessment in three different spatial configurations; 1) S0/N0: speech and noise presented from the front, 2) S_CI_/N_NHE_: speech presented to the CI and noise presented to the NHE and 3) S_0_/N_NHE_: speech presented from the front and noise to the normal hearing ear. All configurations were tested twice, with and without the CI, and block orders were counterbalanced across participants [Távora-Vieira et al., 2015;2016;Wedekind et al., 2018;Wedekind et al., 2020;Wedekind et al., 2021].

### Sound Localisation

Sound localisation was tested using the Auditory Speech Sounds Evaluation Localisation Test. This test was conducted in a sound proof booth and presents a 4000 Hz narrow band noise simultaneously through two loudspeakers that were placed at -60 and 60 degrees from the participant. All stimuli were presented at 60 dB HL at one loud speaker and depnding on the interaural level difference, the other speaker presented at 60, 56, 40 or 30 dB. To create the illusion of a sound source localized somewhere on the azimuth between the two loud speakers, the presentation level from both loud speakers differed to create an interaural level difference of either: -30, -20, -10, -4, 0, +4, +10, +20, +30. The software randomly picks the ILD to present at. This allowed for 13 localisation points to be established, two true speakers and 11 sham speakers. Each speaker was placed in a semicircle at 10 degree intervals in front of the subject, see Figure 1 [Tavora-Vieira et al., 2015].

**Figure 1.**
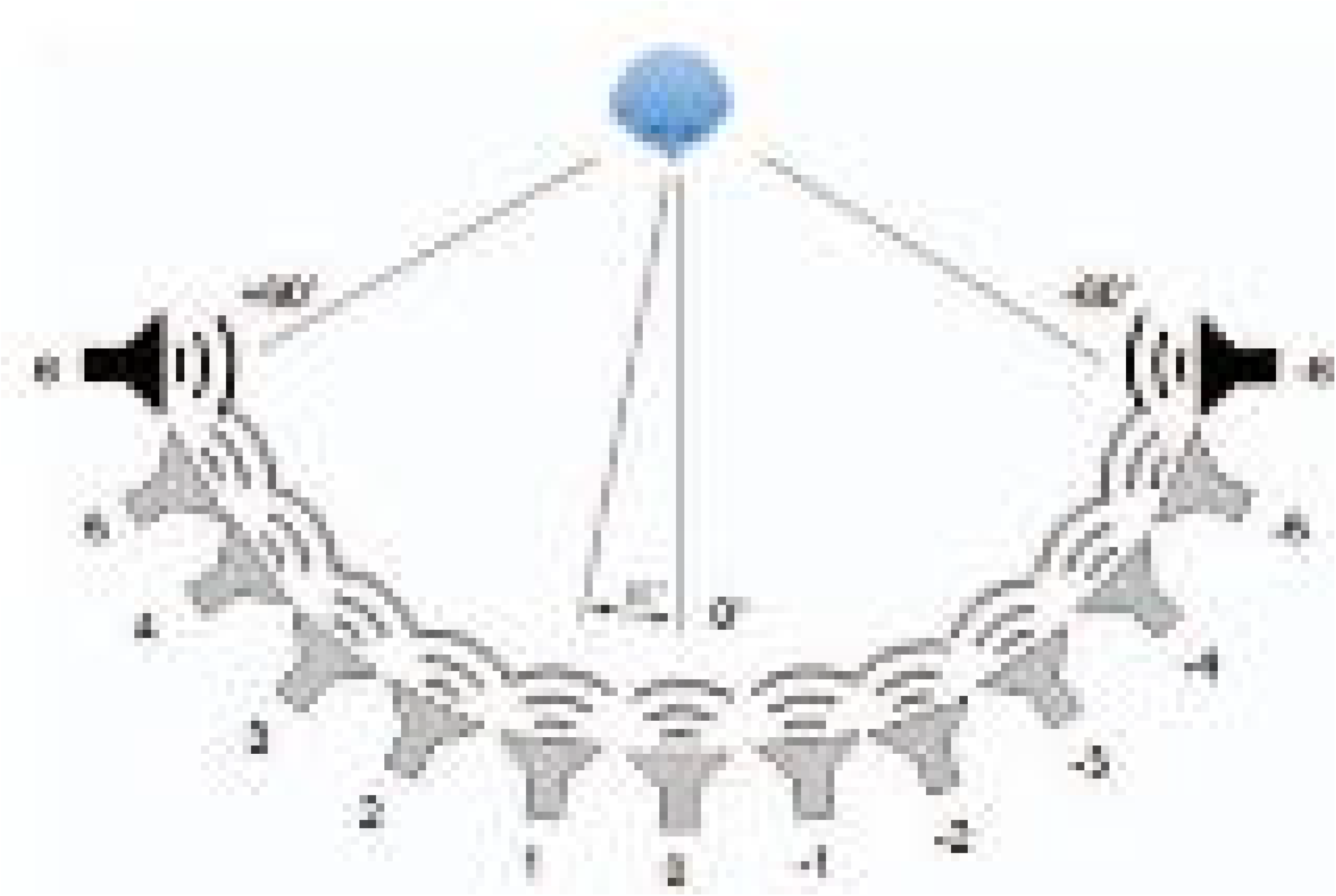
Set up of the localisation test. Participant was seated facing speaker number 0. The black speakers 6 and -6 are the two true speakers and the grey speakers (−5 to 5) are the sham speakers. Each speaker is positioned 10 degrees apart

The thirteen loudspeakers were numbered from -6 to 6. The two real loud speakers were number as -6 and 6 and the 11 sham speakers were numbered from -5 to 5. The participant was required to report which one of the thirteen speakers the sound was coming from. After each response made by the participant, their answer was inputted into the computer software which calculated the median values and root mean square (RMS). A lower RMS indicated better localisation ability.

The total test consisted of 33 items. All narrowband noise presentation locations were randomly selected by a computer software. Stimuly with intensity differences of -30, -20, - 10, 10, 20, 30 dB were presented three times each and stimuli with intensity differences of -4, 0 and 4 dB were presented five times each.

### Oddball Task

This study used a semantic oddball paradigm consisting of odd and even numbers from one to nine that was presented in the presence of background noise. Eight talker background noise wave files were attained from the National Acoustic Laboratories. Speech and noise were presented in free field from two different speakers, 45 degrees azimuth from the subject – with the signal (odd and even numbers) always being presented to the CI side. In all condition participants were instructed to look at a fixation cross presented on a computer monitor 1 metre away from them. This was implemented to reduce eye movement. See Figure 2 for a schematic diagram of the experimental set up for the three conditions.

**Figure 2.**
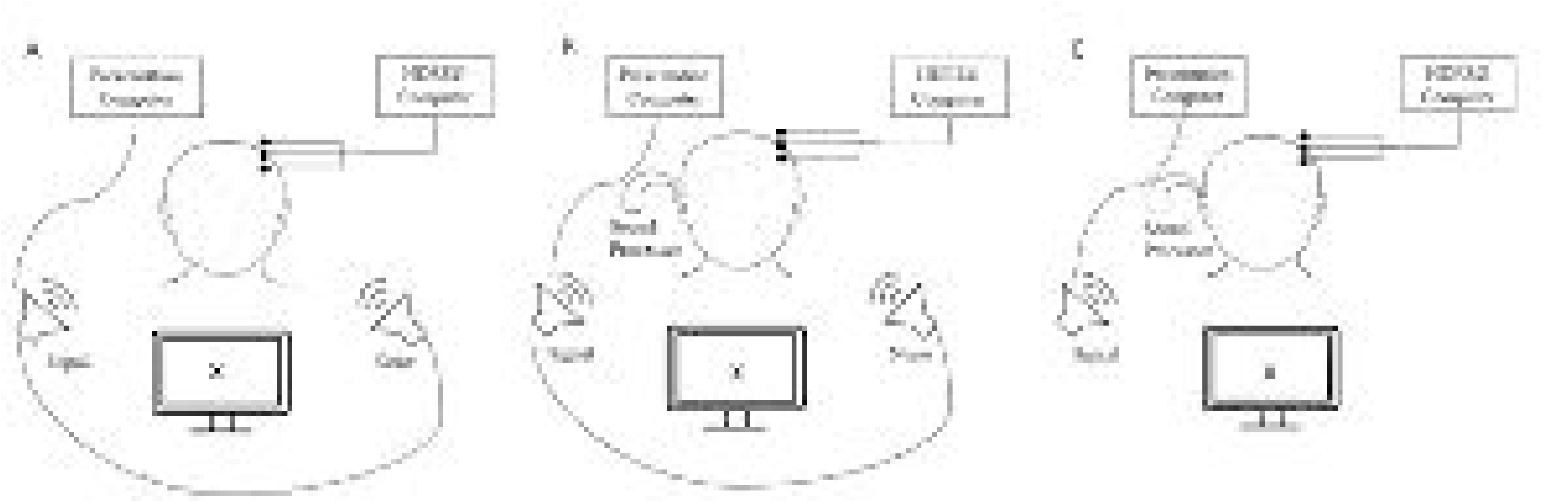
Schematic diagram illustrating the set up of the three experimental conditions. **(a)** depicts CI-Off, **(b)** depicts CI-On and **(c)** depicts Control.

The odd/even oddball paradigm was presented pseudo-randomly such that a target stimulus was presented with a probability of 20% (48 presentations) and a standard stimulus was presented with a probability of 80% (196 presentations). Each stimulus was presented with an inter stimulus interval of 1500ms. In addition, the task order was counter-balanced across participants. The task consisted of odd numbers from one to nine (one, three, five, nine) and even numbers (two, four, six, eight). The number seven was omitted from the odd list as it contains two syllables. These speech files were recorded with the purpose to be used in a telephone-based speech-in-noise test called ‘Telescreen’ [Dillon et al., 2016]. Each recorded number was modified using the software ‘Audacity®’ [Audacity, 1999-2016] so that each number was of an approximate duration of 400ms. Speech babble was presented at 55 dB HL and the numbers were presented at 60 dB HL – resulting in a signal to noise ratio of +5 dB HL.

### Acquisition and Pre-Processing of Electrophysiological Data

Electrophysiology data was continuously recorded for the duration of each condition of the oddball task. The data was acquired using the Micromed™ SD LTM EXPRESS system with Gilat Medical ERP software (Gilat Medical Research & Equipment Ltd, Karkur, Israel). A sampling rate of 1024 Hz with an online low pass-filter of 40 Hz was used to digitise the data. Data was recorded using Ag/AgCl electrode cap (SpesMedica ™ Genova, Italy). The Ag/AgCl electrode cap consisted of 59 electrodes, which were arranged in accordance with the 10-20 system. An additional four electrodes were used to 1) account for myogenic artefact arising due to eyeblinks from an electrode placed under the infraorbital region of the right eye, 2) a reference electrode that was placed on the middle of the chin, 4) a ground electrode placed on the right mastoid. All electrode impedance was kept below 5 kΩ for the duration of the recording.

MATLAB 2020a was used to process the data. A semi-automate procedure was used consisting of functions from the plug-ins EEGLAB [Delorme and Makeig, 2004], PREP pipeline [Bigdely-Shamlo et al., 2015], clean_rawdata() plugin, AMICA [Palmer et al., 2011] and ICLabel plugin [Pion-Tonachini et al., 2019]. The removeTrend() from the PREP pipeline plugin was used to linearly detrend the data using a high pass 1Hz fir filter with a 0.02 step size. The cleanLineNoise() from PREP pipeline plugin was used to remove 50Hz line noise and harmonics up to 500Hz. The pop_clean_rawdata() was used to determine noisy channels. The pop_interp was used to interpolate noisy channels spherically. EEG data was then down sampled to 250 Hz. The data was demeaned and a 30Hz low pass filter was applied using the pop_eegfiltnew(). Filter order equals 100. The clean_asr() was used to correct for artefacts using the artefact subspace reconstruction method. Data was then epoched from -200 to 1000ms relative to stimulus onset. Independent component analysis of the data was conducted using AMICA (2000 iterations) on down sampled data to 100Hz [Palmer et al., 2011]. The number of independent components extracted were adjusted for the data rank. The data was baseline correct was to the pre-stimulus interval (−200 to 0 ms). Trials with activity exceeding 100mV were flagged for exclusion for further analysis. SASICA was used to guide the manual rejection of ICA components that were deemed to be too noisy (mean = 22 components removed).

### Measurement of Event Related Potentials

We measured amplitude of N2N4 and P3b ERP components by calculating the area of standard-target effects on ‘target-minus-standard’ ERP difference waveforms (Fig. 3). These measurements were conducted at the trial-level by subtracting the individual averaged standard ERP of each condition from each individual target trial of the corresponding condition. We measured N2N4 at FCz and P3b at Pz, corresponding to the site where the size of standard-target N2N4 and P3b effects were most prominent. Given the temporally distributed nature of each difference ERP, we used broad time windows (300-800 ms for N2N4 and 500-950 ms for P3b) to capture each component and excluded positive areas for N2N4 and negative areas for P3b. The same time window were used for all three hearing conditions. The latency of the N2N4 and P3b were estimated using the 50% area latency method.

**Figure 3.**
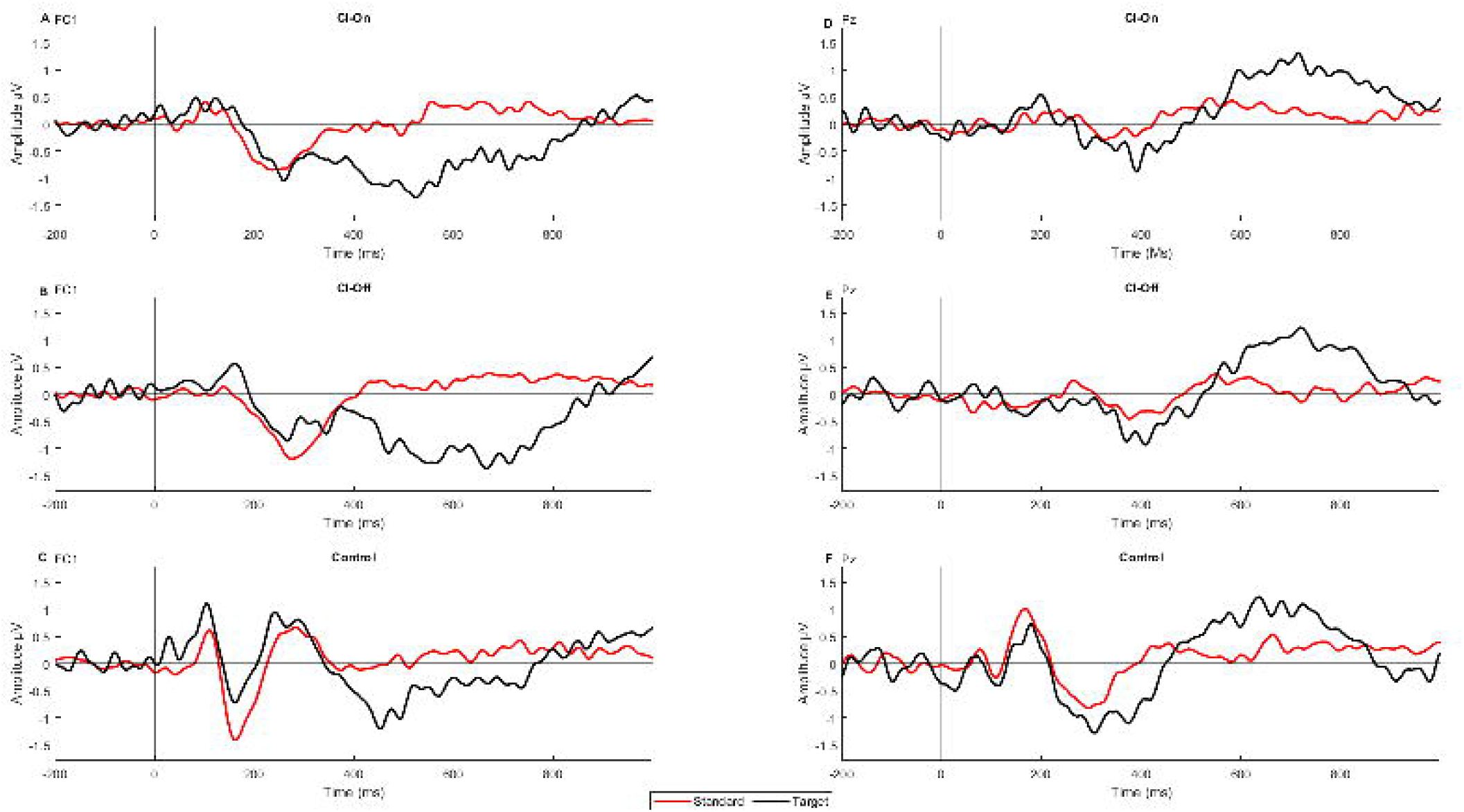
Grand mean ERP waveforms for each stimulus (Standard, Target) and presentation conditions (CI-On, CI-Off, Control). Panel **(a), (b), (c)** depict grand mean waveforms recorded from frontocentral electrode FC1 and panel **(d), (e), (f)** depict grand mean waveforms recorded from parietally distributed electrode Pz.

### Behavioural Data

We examined task performance by measuring RT and RT variability (standard deviation of RTs within each condition), and target accuracy. RTs exceeding than 1500 ms were excluded from further analysis. Target accuracy was calculated as the proportion of target trials that were responded within the accepted window. Subjective listening effort was also measured after the completion of each condition. This was measured by using a seven-point scale where 1 indicated ‘No Effort’ and 7 indicated ‘Extreme Effort’. Participants verbally indicated which number corresponded to their perceived listening effort [Luts et al., 2010;Holube et al., 2016].

### Statistical Analysis

All statistical analysis were conducted using R statistics and R Studio software [R, 2013]. We conducted linear mixed model analysis using the ‘lme’ function from the ‘nlme’ package [Pinheiro et al., 2022]. Localisation, speech in noise, RT, subjective listening effort and Target Accuracy, we included condition (CI-Off, CI-On and Control) as a fixed effect, and intercepts for participants were modelled as a random effect. For electrophysiological measures (N2N4, P3b) we also included Trial-Type as an additional fixed effect, interaction with the effect of Ear.

Results were analysed using the ‘anova’ function and presented as F-values. Follow-up pairwise comparison were conducted using the ‘emmeans’ and ‘contrast’ function from the ‘emmeans’ package [Lenth et al., 2020]. Pairwise results were presented as t-rations (mean difference estimate divided by standard error) and p-values for multiple comparisons were corrected using the ‘Holm’ method. The function ‘emmeans’ was also used to plot values and error bars for figures presented in the results.

## Results

### Localisation and Speech in Noise

For localisation, the linear mixed model analysis revealed a significant effect of CI (*F*(1,49) = 102.79, *p* < .0001*), indicating an improvement in sound localisation ability with the CI-on compared to CI-off (*M*(*SD*) = 23.6(11.1) vs. 50.0(18.0) degrees; *Est. Mean Diff*. (*SE*) = 26.4(2.6) degrees) (Fig. 4A).

**Figure 4.**
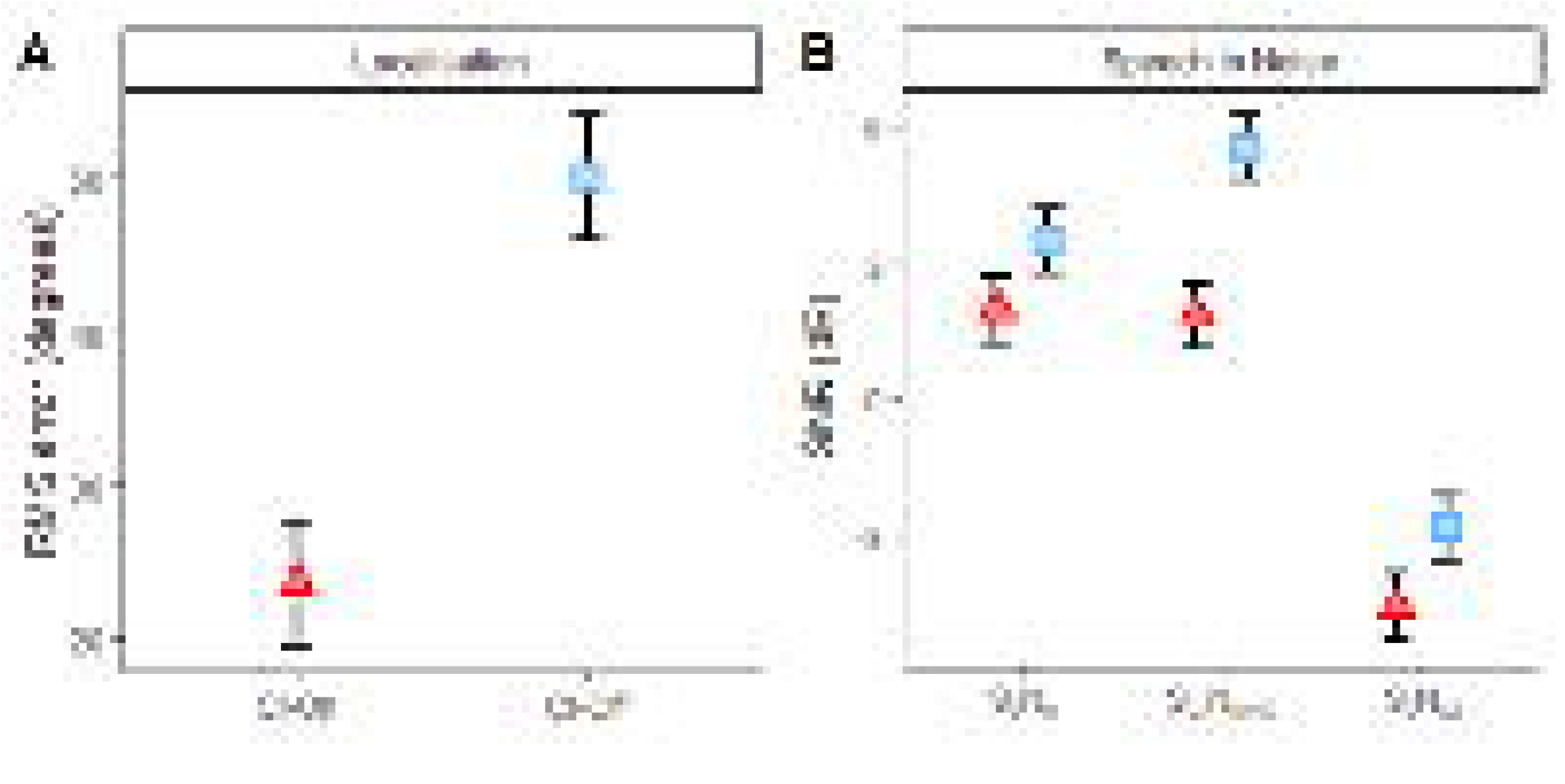
**(a)** depicts group means with within subject standard error bars for speech in noise intelligibility using the Bamford-Kowol-Bench Speech-In-Noise Test. Test was conducted in three spatial configurations, with and without the CI; S0N0 – speech and noise from front, Sci/Nhe – speech from CI side, noise from NHE side and S0/Nci – speech from front, noise from CI side. **(b)** Sound localisation test results with and without the CI. A lower RMS indicates better sound localisation ability.

For Speech in Noise, the linear mixed model analysis revealed significant main effects of CI (*F*(1,45) = 22.74, *p* < .0001*) and sound presentation (*F*(2,45) = 90.94, *p* < .0001*) but the two-way interaction was not statistically significant (*F*(2,45) = 2.05, *p* = .141). Pairwise contrasts between CI-on and CI-off for each configuration revealed there was a significant improvement S_CI_N_NHE_ (*t-ratio*(45) = 4.40, *p* = .0002*, *Est. mean diff*. (*SE*) = 3.70 (0.84)) but not in S_0_N_0_ (*t-ratio*(45) = 2.08, *p* = .087, *Est. mean diff*. (*SE*) = 1.75 (0.84))and S_0_N_CI_ conditions (*t-ratio*(45) = 1.78, *p* = .087, *Est. mean diff*. (*SE*) = 1.50 (0.84)) (Fig. 4B).

### Reaction Time, Target Accuracy and Subjective Effort

For reaction time, the linear mixed model analysis revealed a significant effect of task condition (*F*(2,1624) = 29.34, *p* < .0001*). RTs were shortest in Control condition (no-noise with CI-on, M(SD) = 784(143) ms) followed by CI-on (807(126) ms) then CI-off with noise (850(161) ms) (Fig. 5A). Differences in RT between task conditions were all statistically significant (Control vs. CI-on: *t-ratio*(1624) = 3.08, *p* = .0021*, *Est. mean diff*. (*SE*) = 24.4(7.9) ms; Control vs. CI-off: *t-ratio*(1624) = -7.62, *p* < .0001*, *Est. mean diff*. (*SE*) = 60.6(7.95) ms, CI-on vs. CI-off: *t-ratio*(1624) = 4.54, *p* < .0001*, *Est. mean diff*. (*SE*) = 36.2(7.97) ms).

**Figure 5.**
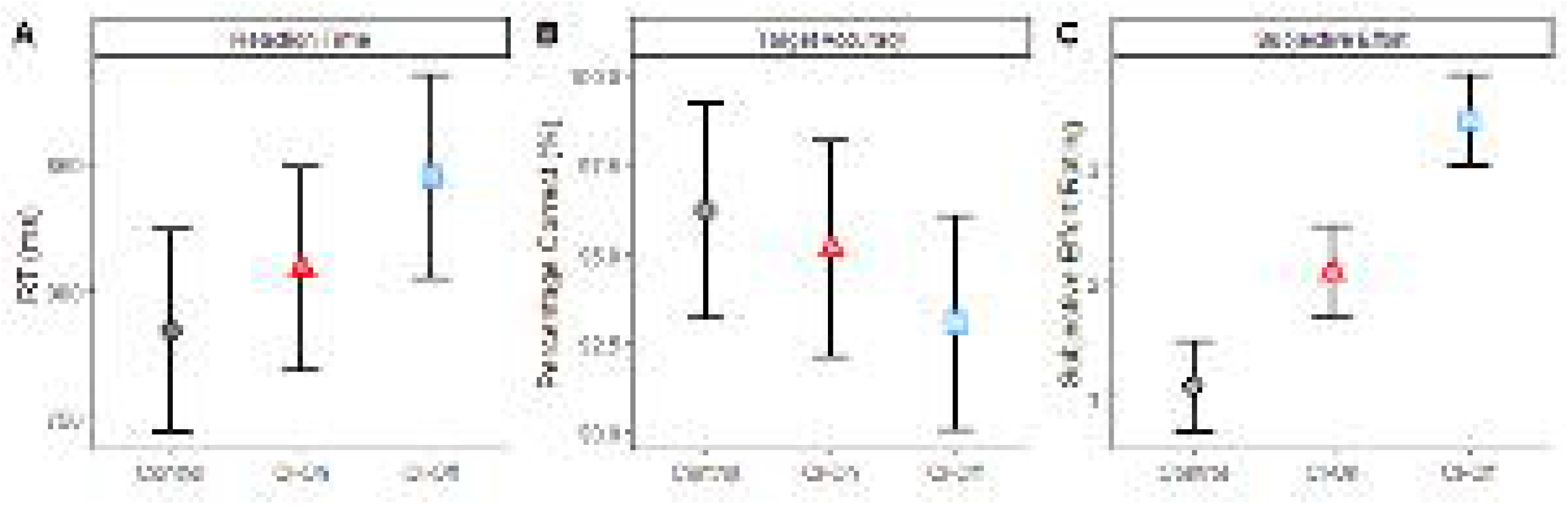
Grand mean estimates with error bars depicting the standard error of the mean. **(a)** depicts the reaction time, **(b)** depicts the target accuracy and **(c)** depicts the subjective listening effort.

Target accuracy (Fig. 5B) exceeded 90% on all task condition with the lowest means for CI-off and the highest means for Control (Control: M(SD) = 96.18(8.42) %; CI-on: 95.14(9.91) %; CI-off: 93.06(12.60) %). However, the linear mixed model analysis indicated that differences in accuracy between task conditions were not statistically significant (*F*(2,22) = 2.66, *p* = .0927).

Subjective listening effort (Fig. 5C) measured after the completion of each condition revealed that participants perceived the CI-Off condition required the greatest listening effort, followed by CI-On and then Control condition (Control: M(SD) = 1.08(1.38), CI-on = 2.08(1.62), CI-off = 3.42(1.08)). The linear mixed model analysis revealed a statistically significant effect of task condition (*F*(2,22) = 22.93, *p* < .0001*). Follow-up pairwise comparisons showed that differences in subjective effort ratings between all task conditions were statistically significant (Control vs. CI-on: *t-ratio*(22) = 2.89, *p* = .0221*, *Est. mean diff*. (*SE*) = 1(0.25); Control vs. CI-off: *t-ratio*(22) = 6.75, *p* < .0001*, *Est. mean diff*. (*SE*) = 2.33(0.25); *t-ratio*(22) = 3.86, *p* = .0024*, *Est. mean diff*. (*SE*) = 1.33(0.35)).

### N2N4 area amplitude and 50% area latency

N2N4 area amplitude was calculated using difference waveforms using a frontocentral electrode (Fig. 6). The linear mixed model analysis showed that there was no statistically significant difference in area amplitude between task condition (*F*(2,1614) = 1.24, *p* = .289). However, there was a significant main effect of task condition for latency (*F*(2,1624) = 5.4, *p* = .0045*). As depicted in Figure 4, mean latency was shortest for Control (M(SD) = 520(33) ms), followed by CI-On (535(26) ms) and CI-Off (552(42) ms). Follow-up pairwise comparisons revealed that differences in area latency between Control and CI-off were statistically significant (*t-ratio*(1624) = 3.28, *p* = .0032*, *Est. mean diff*. (*SE*) = 32.8(10) ms), but not for Control vs. CI-On (*t-ratio*(1624) = 1.36, *p* = .175, *Est. mean diff*. (*SE*) = 13.5(9.94) ms) and CI-On vs. CI-Off (*t-ratio*(1624) = 1.93, *p* = .1079, *Est. mean diff*. (*SE*) = 19.3(10.03) ms).

**Figure 6.**
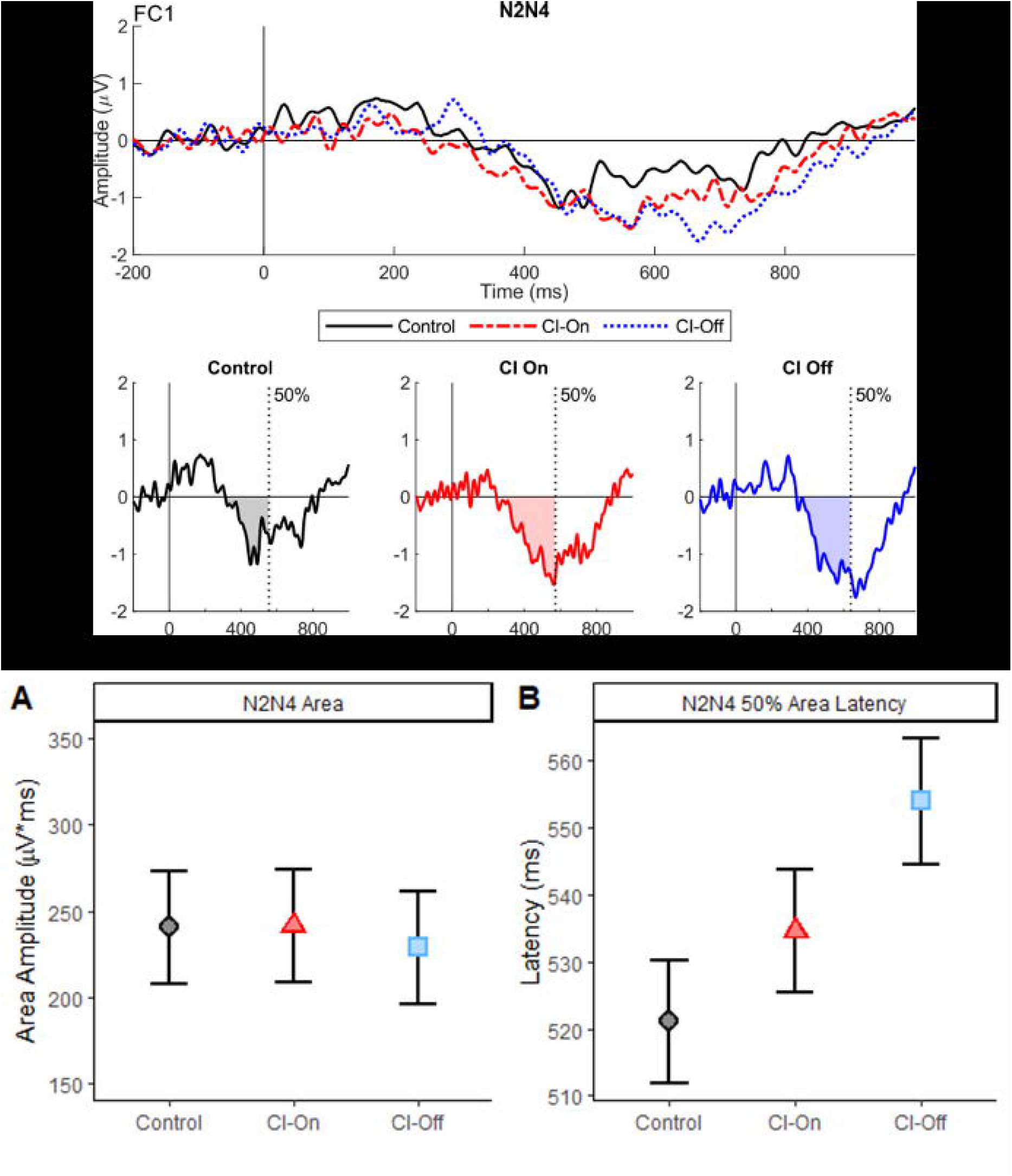
ERPs measured from a frontocentral electrode (FCz). **(a)** depicts the difference waveform (target minus standard) for all three testing conditions. The grey highlighted region indicates the time window used to measure the N2N4 (300 to 800 ms). **(b) & (c)** mean area and latency reflecting the N2N4 measured in all three conditions, respectively.

### P3b area amplitude and 50% area latency

Difference wave forms (target minus standard) was used to calculate the P3b area using a parietal electrode (Fig. 7). Linear mixed model analysis revealed a significant main effect of task condition (*F*(2,1612) = 4.34, *p* = .0132*). Follow-up pairwise comparisons showed that P3b area amplitude was significantly greater for Control compared to CI-on (*t-ratio*(1612) = 2.43, *p* = .0305*, *Est. mean diff*. (*SE*) = 21.30 (8.77) μV*ms) and CI-off (*t-ratio*(1612) = 2.65, *p* = .0242*, *Est. mean diff*. (*SE*) = 23.39(8.82) μV*ms), but not between CI-on vs. CI-off (*t-ratio*(1612) = 0.24, *p* = .8127, *Est. mean diff*. (*SE*) = 2.09(8.84) μV*ms).

**Figure 7.**
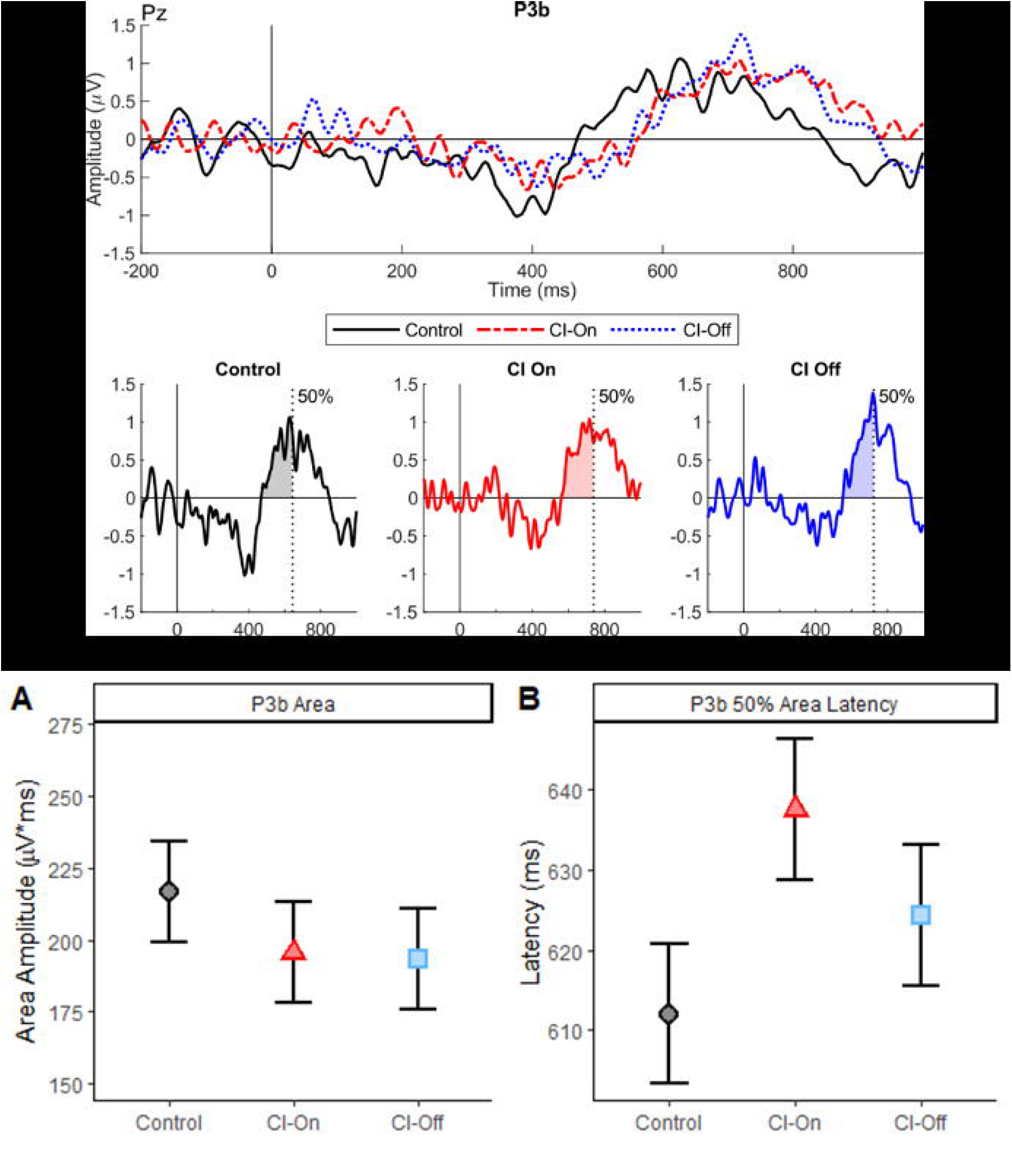
ERPs measured from a posterior electrode (Pz). **(a)** depicts the difference waveform (target minus standard) for all three testing conditions. The grey highlighted region indicates the time window used to measure the P3b (500 to 950 ms). **(b) & (c)** mean area and latency reflecting the P3b measured in all three conditions, respectively.

Looking at P3b 50% area latency, the main effect of task condition was approaching statistical significance (*F*(2,1612) = 2.90, *p* = .056). Follow-up pairwise comparisons, revealed P3b area latency was significantly shorter for Control compared to CI-On (*t-ratio*(1612) = 2.41, *p* = .048*, *Est. mean diff*. (SE) = 25.5 (10.6) ms) but differences between Control vs. CI-off (*t-ratio*(1612) = 1.15, *p* = .249, *Est. mean diff*. (*SE*) = 12.3 (10.6) ms), and CI-on vs. CI-off (*t-ratio*(1612) = 1.24, *p* = .249, *Est. mean diff*. (*SE*) = 13.2(10.7) ms) were not statistically significant.

## Discussion

In the present study, we examined the neural processing of of words presented during background noise in SSD CI users. In particular, we focused on understanding how the CI impacts the ability to discriminate odd/even numbers by comparing ERP results obtained with and without the CI, and assessed the effect of noise by contrasting the results with a no-noise (Control) condition. We also characterised functional hearing ability, by measuring sound localisation and speech in noise intelligibility with and without the CI. In the functional hearing task, we identified a significant improvement in both test when the CI was switched on. In the semantic oddball task, the best performance was observed during the no-noise condition, but under noisy conditions, participants performed better during CI-On compared CI-Off, as indicated by faster RT, higher target accuracy and a lower subjective listening effort rating. For ERPs, we observed an effect of condition on N2N4 latency (Control < CI-On < CI-Off), and P3b amplitude (Control > CI-On/Off) and latency (Control < CI-On/Off).

### Functional Hearing: Speech In Noise and Sound Localisation

Functional improvement with CI was observed during the speech in noise test. The improvement was most prominent when the speech signal was directed at the CI-side (S_CI_N_NHE_). Smaller CI-related improvements were also observed in S_0_N_0_ and S_0_N_CI_ configurations, however these did not reach the threshold for statistical significance (*p* = 0.87). These smaller effects highlight the dominance of the NHE in S_0_N_0_ [Van de Heyning et al., 2008;Arndt et al., 2011;Dorbeau et al., 2018] and that the CI is not deteimental to speech intelligibility when noise is coming from the CI side [Wedekind et al., 2021]. Likewise, a statistically significant improvement in sound localisation was seen with the CI on, consistent with previous studies [Vermeire and Van de Heyning, 2009;Firszt et al., 2012;Távora-Vieira et al., 2015;Wedekind et al., 2021]. Collectively, the reuslts demonstrate that the CI is capable of restoring binaural hearing in SSD patients.

### Semantic Oddball Task: Task Performance

In line with the functional hearing results, RTs to target stimuli in the Oddball task were faster during CI-On compared to CI-Off, indicating that the addition signal from the CI significantly facilitated the ability to process and identify target words. RTs to the Control condition (no-noise with CI) was significantly shorter than both CI-On and CI-Off. This performance increase was also accompanied by participant reports that less effort was required to perform the task with CI-On compared to CI-Off, demonstrating that the use of CI had a noticable benefit on perceieved task difficulty. Similarly, subjective effort ratings to the control condition was significantly lower than CI-On and CI-Off. Although reactions were slowest and the task perceived most effortful during CI-Off, participants were highly accurate across all conditions (< 90%). While the main effect of condition was not statistically significant, inspection of mean values indicated that CI-Off was lowest (93%), followed by CI-On (95.1%) and Control (96.2%), which is in line with the RT and subjective effort ratings.

Collectively, these behavioural results demonstrated that (1) the use of background noise had a significant impact on objective performance as well as subjective perceptions of task difficulty – but participants were able to successfully complete the task descpite the noise – and (2) that use of background noise allowed us to show a measurable improvement in bilateral hearing (CI-On) comapred to hearing with the NHE alone (CI-Off).

### Semantic Oddbal Task: Neural Responses

With respect to neural processing, we identified that with the CI-Off, N2N4 latency was signficiantly delayed compared to the Control condition. The delay in N2N4 during the more difficult (CI-Off) condition is consistent with previous within-group observations, such as Almeqbel and McMahon (2015) who examined neural responses to speech-tokens in background noise using a passive task with young children. They reported delayed N2N4 latencies in with lower (−10 dB) compared to higher SNR conditions (+20 dB). Our results are also consistent with Finke et al. (2016) who reported delayed N2N4 latencies in CI users comapred to a normal hearing control group. One interpretation for the delay is that it reflects the increased effort in accessing lexical information during adverse listening conditions (Finke et al. 2016). Alternatively, the delay could also reflect increased effort needed to resolve lower-level uncertainty in the sound signal whereby previous studies have attributed this uncertainty to the CI (Cope et al. 2017; Obleser et al. 2007), but in our study we attribute this uncertianty to the background noise. Although our task required word discrimination and our N2N4 results could be interpreted in the context of retrieving word meanings from the mental lexicon, we cannot rule out the possibility that our results may be driven by more general lower-level uncertainties. Nevertheless, our N2N4 findings suggest that the higher-order cognitive processes involved in evaluating the simple words requires more time under noisy conditions, without the CI.

With respect to P3b area latency we observed that the Control condition had a signficantly shorter latencies when compared with the CI-On. However, the difference between CI-Off and CI-On/Control were not statistically significant. With respect to amplitude, a similar pattern was observed, where P3b amplitudes were more positive during control compared to CI-On, but no differences was observed between CI-On and CI-Off.

Focusing on P3b area latency we observed that the no noise Control condition had a signficantly shorter latency when compared with noise conditions (CI-On and CI-Off) [Polich, 2007]. Additionally we identified signficantly shorter RT and lower percieved subjective listening effort results for the Control condition. Taken together the P3b area latency and behavioural results indicate that in the absence of noise, SSD CI users are able to discriminate and evluate auditory stimuli quicker relative to in environments with noise. Despite the lack of statistically signficant differences in P3b area between CI conditions, we did identify that the P3b area was largest in the control condition which was identified to be the easiest conditions indicated by smallest RT and subjective listening scores. The larger P3b area for the Control condition suggest that in the absence of noise, evaluation of auditory stimuli is easier when compared to situations in noisy environments [Polich, 2007].

The P3b area and latency findings are surprising given that functional assessments indicated that with the CI, SSD CI users record improvements in speech in noise intelligibilty with the CI. We would have expected this improvemnet in functional assessment, along with the behavioural results (RT and subjective listeining effort), to result in larger P3b areas and earlier latency being recroded for CI-On when compared with CI-Off, but we did not observe this between CI-On and CI-Off. This finding alludes to the possibility that EEG may not be sensitive enough to detect within subject differences. The clear differences observed in the behavioural data and the lack of differences in the EEG data suggest that there are limitations with using EEG as a measure of cognitive processing.

### Reliability of ERPs as measures of cognitive processing

The lack of consistency in the explanation of N2N4/P3b responses in the present and previous studies highlights that using N2N4/P3b to measure cognitive effort may be more complex and subjected to large amounts of varibility. Finke et al identified that the N2N4 amplitude recorded from directly stimulating the CI was larger than when recording directly from the NHE and attributed this increase in N2N4 during direct connect of CI to lexical processing. This larger N2N4 reflected greater effort to match the sound with the mental lexicon, showing support for the conflict monitoring hypothesis [Finke et al., 2016]. Conversely, in previous work conducted by our lab, we identified that a smaller N2N4 was recorded from directly stimulating the CI in comparison to the NHE [Wedekind et al., 2021;Voola et al., 2022]. Wedekind et al (2021) used an auditory oddball paradigm consisting of pure tones (1kHz and 2kHz) identitfying that compared to the NHE, the CI showed a smaller N2 but the P3b was similar between the NHE and CI. However, given the simplicity of the pure tone oddball task, the paper highlighted the possibility that stimulus differences were mainly discriminated by early discrimination process reflected by the N2 and deeper evaluation of stimulus was not needed [Wedekind et al., 2021].

To build on the findings of Wedekind et al. (2021), our lab conducted a follow-up study whereby SSD CI users had to discriminate between odd and even numbers, comparing both the NHE vs CI (both via direct stimulation) and also in free field, with and without the CI. We identified that N2N4 and P3b were both similar between the NHE and CI, eventhough the behavioural data indicated that evaluation of auditory stimuli from the CI was significantly slower in RT when compared to the NHE. For free field, we observed similar N2N4, P3b and RT results between CI-On and CI-Off, which suggested that the NHE was dominating the response. This rationale was developed due to the stimuli used in the study not containing any binaural cues. As such the current study was implemented noise using an audtory oddball tasks.

With the ambition to create a task that is more complex, the current and past study’s [Finke et al., 2017] have added noise to the auditory oddball task with the aim that increasing task complexity will reveal more about the higher order process of SSD CI users. Finke et al. comapred CI users with normal hearing controls, evaluating their higher order proccessing of speech using german two syllable words in noise. Overall the current study and Finke et al. identified that in the no noise condition, SSD CI participants performed better behaviouraly (shorter RTs) than compared to the tasks that had noise. However, the N2N4 and P3b results were from both studies showed mixed results. This is evident in the P3b area findings in Finke et al (2017) who identified that in the most complex task (modualted noise) P3b area was larger when compared to the no noise condition. Conversley, in the current study we observed that the most complex condition (CI-Off) elicited the smallest P3b area, attributing this effect to stimuli being more difficult to distinguish. These inconsistencies in EEG data between studies highlights the large variability with using N2N4 and P3b to measure cognitive ability. Additionally, implementing noise into an auditory oddball task may not be the answer to be able to gain a deeper understanding of cognitive ability of SSD CI users. This is highlighted by the fact that early ERPs (N1-P2) which are thought to have downstream effects on later ERPs (N2N4 & P3b), cannot be identified in the waveforms generated from noisy conditions, but can be seen in the no-noise condition (Fig. 3). The absence of clear early ERPs highlight that the adding more noise to the auditory oddball paradigm will only result in the waveforms generated being harder to interpret and being able to compare with past research from no-noise studies. To overcome these issues with N2N4 and P3b measurements, future studies should look to employ alternative measures such as pupillometry which has been shown in past literature to be a good measure of cognitive effort [Piquado et al., 2010;López-Ornat et al., 2018].

## Conclusion

In the present study, we identified signficant differences in RT, subjective listening effort, both indicating that the Control condition was both objectively and subjectively the easiest condition. Despite significant differences in RT, the neural responses (N2N4 and P3b) did not follow the same trend for all three conditions. The lack of consistency between the behavioural and neural repsonses highlights the variability in using N2N4 and P3b to measure cognitive effort. This was emphasied by previous stuides employing different explanation for N2N4 and P3b effects. This highlights the need for caution to be taken when designing auditory oddball tasks with CI patients in future studies. By using other forms of measures for cognitive ability (such as pupillometry) in speech in noise task with SSD CI users, this knowledge could potentially guide implantation candidacy guidelines and management rehabilitation protocols.

## Acknowledgments

National Acoustics Laboratories provided the eight-talker speech babble.

## Statement Ethics

Ethics approval was obtained from the South Metropolitan Health Ethics Committee (reference number: 335). Participants have given their written informed consent to participate in this study.

## Author Contributions

**Marcus Voola:** Drafting, design, data collection, interpretation, final approval. **Andre Wedekind:** Drafting, design, interpretation. **An Nguyen:** Drafting, design, analysis, interpretation. **Welber Marinovic:** Drafting, final approval, interpretation. **Gunesh Rajan:** Interpretation, drafting, final approval. **Dayse Tavora-Vieira:** Interpretation, drafting, final approval, design

## Data Availability Statement

Data can be made available upon request sent to the corresponding author.

## Legends

**Table 1:** Demographic information of participants age, gender, duration of deafness, cause of deafness (ISSNHL = Idiopathic Sudden Sensorineural Hearing Loss, MD = Meniere’s Disease), side of implant, pure tone average (PTA), inserted electrode type and experience with the CI.

## References

Arndt S, Aschendorff A, Laszig R, Beck R, Schild C, Kroeger S, et al. Comparison of Pseudobinaural Hearing to Real Binaural Hearing Rehabilitation After Cochlear Implantation in Patients With Unilateral Deafness and Tinnitus. Otology & Neurotology. 2011;32(1):39–47.

Audacity. Audacity(R). 1999-2016. p. The name Audacity(R) is a registered trademark of Dominic Mazzoni.

Bench J, Kowal Å, Bamford J. The Bkb (Bamford-Kowal-Bench) Sentence Lists for Partially-Hearing Children. British Journal of Audiology. 1979 1979/01/01;13(3):108–12.

Bigdely-Shamlo N, Mullen T, Kothe C, Su K-M, Robbins KA. The PREP pipeline: standardized preprocessing for large-scale EEG analysis. Frontiers in Neuroinformatics. 2015 2015-June-18;9(16).

Comerchero MD, Polich J. P3a and P3b from typical auditory and visual stimuli. Clinical neurophysiology : official journal of the International Federation of Clinical Neurophysiology. 1999 Jan;110(1):24–30.

Deacon D, Breton F, Ritter W, Vaughan HG. The Relationship Between N2 and N400: Scalp Distribution, Stimulus Probability, and Task Relevance. Psychophysiology. 1991;28(2):185–200.

Delorme, Makeig. EEGLAB: an open source toolbox for analysis of single-trial EEG dynamics including independent component analysis. Journal of neuroscience methods. 2004 Mar 15;134(1):9–21.

Dillon H, Beach EF, Seymour J, Carter L, Golding M. Development of Telescreen: a telephone-based speech-in-noise hearing screening test with a novel masking noise and scoring procedure. Int J Audiol. 2016 Aug;55(8):463–71.

Dorbeau C, Galvin J, Fu QJ, Legris E, Marx M, Bakhos D. Binaural Perception in Single-Sided Deaf Cochlear Implant Users with Unrestricted or Restricted Acoustic Hearing in the Non-Implanted Ear. Audiology & neuro-otology. 2018;23(3):187–97.

Drennan WR, Rubinstein JT. Music perception in cochlear implant users and its relationship with psychophysical capabilities. Journal of rehabilitation research and development. 2008;45(5):779–89.

Finke M, Büchner A, Ruigendijk E, Meyer M, Sandmann P. On the relationship between auditory cognition and speech intelligibility in cochlear implant users: An ERP study. Neuropsychologia. 2016 2016/07/01/;87:169–81.

Finke M, Sandmann P, Bönitz H, Kral A, Büchner A. Consequences of Stimulus Type on Higher-Order Processing in Single-Sided Deaf Cochlear Implant Users. Audiology and Neurotology. 2017;21(5):305–15.

Firszt JB, Holden LK, Reeder RM, Waltzman SB, Arndt S. Auditory abilities after cochlear implantation in adults with unilateral deafness: a pilot study. Otology & neurotology : official publication of the American Otological Society, American Neurotology Society [and] European Academy of Otology and Neurotology. 2012 Oct;33(8):1339–46.

Friedmann DR, Ahmed OH, McMenomey SO, Shapiro WH, Waltzman SB, Roland JT, Jr. Single-sided Deafness Cochlear Implantation: Candidacy, Evaluation, and Outcomes in Children and Adults. Otology & neurotology : official publication of the American Otological Society, American Neurotology Society [and] European Academy of Otology and Neurotology. 2016 Feb;37(2):e154–60.

Galvin JJI, Fu Q-J, Wilkinson EP, Mills D, Hagan SC, Lupo JE, et al. Benefits of Cochlear Implantation for Single-Sided Deafness: Data From the House Clinic-University of Southern California- University of California, Los Angeles Clinical Trial. Ear and hearing. 2019;40(4):766–81.

Holube I, Haeder K, Imbery C, Weber R. Subjective Listening Effort and Electrodermal Activity in Listening Situations with Reverberation and Noise. Trends Hear. 2016 Oct 3;20.

Johnson R. The amplitude of the P300 component of the event-related potential: Review and synthesis. Adv Psychophysiol. 1988 01/01;3.

Körtje M, Eichenauer A, Stöver T, Baumann U, Weissgerber T. Impact of Reverberation on Speech Perception and Sound Localization Accuracy in Cochlear Implant Users With Single-Sided Deafness. Otology & neurotology : official publication of the American Otological Society, American Neurotology Society [and] European Academy of Otology and Neurotology. 2022 Jan 1;43(1):e30–e37.

Lau, Phillips, Poeppel. A cortical network for semantics: (de)constructing the N400. Nature Reviews Neuroscience. 2008 2008/12/01;9(12):920–33.

Lenth R, Singman H, Love J, Buerknerm P, Herve M. emmeans: Estimated Marginal Means, aka Least-Squares Means. 2020.

Light GA, Williams LE, Minow F, Sprock J, Rissling A, Sharp R, et al. Electroencephalography (EEG) and event-related potentials (ERPs) with human participants. Curr Protoc Neurosci. 2010 Jul;Chapter 6:Unit 6.25.1-24.

López-Ornat S, Karousou A, Gallego C, Martín L, Camero R. Pupillary Measures of the Cognitive Effort in Auditory Novel Word Processing and Short-Term Retention. Frontiers in Psychology. 2018 2018-November-27;9.

Luts H, Eneman K, Wouters J, Schulte M, Vormann M, Buechler M, et al. Multicenter evaluation of signal enhancement algorithms for hearing aids. The Journal of the Acoustical Society of America. 2010 Mar;127(3):1491–505.

Ma N, Morris S, Kitterick PT. Benefits to Speech Perception in Noise From the Binaural Integration of Electric and Acoustic Signals in Simulated Unilateral Deafness. Ear and hearing. 2016;37(3):248–59.

Näätänen, Picton. N2 and automatic versus controlled processes. Electroencephalography and clinical neurophysiology Supplement. 1986;38:169–86.

Palmer, Kreutz-Delgado, Makeig. AMICA : An Adaptive Mixture of Independent Component Analyzers with Shared Components. 2011.

Pinheiro J, Bates D, DebRoy S. nlme: Linear and Nonlinear Mixed Effects Models. 2022.

Pion-Tonachini L, Kreutz-Delgado K, Makeig S. The ICLabel dataset of electroencephalographic (EEG) independent component (IC) features. Data in Brief. 2019 2019/08/01/;25:104101.

Piquado T, Isaacowitz D, Wingfield A. Pupillometry as a measure of cognitive effort in younger and older adults. Psychophysiology. 2010 May 1;47(3):560–9.

Polich. N400s from sentences, semantic categories, number and letter strings? Bulletin of the Psychonomic Society. 1985 1985/04/01;23(4):361–64.

Polich. Attention, probability, and task demands as determinants of P300 latency from auditory stimuli. Electroencephalography and clinical neurophysiology. 1986;63(3):251–59.

Polich. Updating P300: an integrative theory of P3a and P3b. Clinical neurophysiology : official journal of the International Federation of Clinical Neurophysiology. 2007 Oct;118(10):2128–48.

Polich J, Ladish C, Burns T. Normal variation of P300 in children: age, memory span, and head size. International journal of psychophysiology : official journal of the International Organization of Psychophysiology. 1990 Oct;9(3):237–48.

R TR. A language and environment for statistical computing. Vienna, Austria2013.

Soshi T, Hisanaga S, Kodama N, Kanekama Y, Samejima Y, Yumoto E, et al. Event-related potentials for better speech perception in noise by cochlear implant users. Hear Res. 2014 2014-Oct- ;316:110–21.

Távora-Vieira, Marino R, Acharya A, Rajan GP. The impact of cochlear implantation on speech understanding, subjective hearing performance, and tinnitus perception in patients with unilateral severe to profound hearing loss. Otology & neurotology : official publication of the American Otological Society, American Neurotology Society [and] European Academy of Otology and Neurotology. 2015 Mar;36(3):430–6.

Távora-Vieira, Marino R, Acharya A, Rajan GP. Cochlear implantation in adults with unilateral deafness: A review of the assessment/evaluation protocols. Cochlear Implants Int. 2016 Jul;17(4):184–89.

Tavora-Vieira D, De Ceulaer G, Govaerts PJ, Rajan GP. Cochlear implantation improves localization ability in patients with unilateral deafness. Ear and hearing. 2015 May-Jun;36(3):e93–8.

Van de Heyning P, Vermeire K, Diebl M, Nopp P, Anderson I, De Ridder D. Incapacitating unilateral tinnitus in single-sided deafness treated by cochlear implantation. The Annals of otology, rhinology, and laryngology. 2008 Sep;117(9):645–52.

van den Brink D, Brown CM, Hagoort P. Electrophysiological evidence for early contextual influences during spoken-word recognition: N200 versus N400 effects. Journal of cognitive neuroscience. 2001 Oct 1;13(7):967–85.

Verleger R. On the utility of P3 latency as an index of mental chronometry. Psychophysiology. 1997;34(2):131–56.

Verleger R, Baur N, Metzner MF, Śmigasiewicz K. The hard oddball: Effects of difficult response selection on stimulus-related P3 and on response-related negative potentials. Psychophysiology. 2014;51(11):1089–100.

Vermeire K, Van de Heyning P. Binaural hearing after cochlear implantation in subjects with unilateral sensorineural deafness and tinnitus. Audiology & neuro-otology. 2009;14(3):163–71.

Voola M, Nguyen A, Wedekind A, Marinovic W, Rajan G, Tavora-Vieira D. A Study of Event-Related Potentials during Monaural and Bilateral Hearing in Single Sided Deaf Cochlear Implant Users. bioRxiv. 2022:2022.06.14.495873.

Voola M, Távora-Viera D. Quality of Life handicap measured in patients with profound unilateral or bilateral deafness. Tasman Medical Journal. 2021;3(1).

Wedekind, Rajan G, Van Dun B, Távora-Vieira D. Restoration of cortical symmetry and binaural function: Cortical auditory evoked responses in adult cochlear implant users with single sided deafness. PLoS ONE. 2020;15(1):e0227371.

Wedekind, Tavora-Vieira, Rajan. Cortical auditory evoked responses in cochlear implant users with early-onset single-sided deafness: indicators of the development of bilateral auditory pathways. Neuroreport. 2018 Mar 21;29(5):408–16.

Wedekind, Távora-Vieira D, Nguyen AT, Marinovic W, Rajan GP. Cochlear implants in single-sided deaf recipients: Near normal higher-order processing. Clinical Neurophysiology. 2021 2021/02/01/;132(2):449–56.

Williges B, Wesarg T, Jung L, Geven LI, Radeloff A, Jürgens T. Spatial Speech-in-Noise Performance in Bimodal and Single-Sided Deaf Cochlear Implant Users. Trends Hear. 2019 Jan-Dec;23:2331216519858311.

